# ER membranes exhibit phase behavior at sites of organelle contact

**DOI:** 10.1101/707505

**Authors:** Christopher King, Prabuddha Sengupta, Arnold Seo, Jennifer Lippincott-Schwartz

## Abstract

The plasma membrane of cells exhibits phase behavior that allows transient concentration of specific proteins and lipids, giving rise to functionally dynamic and diverse nanoscopic domains. This phase behavior is observable in giant plasma membrane-derived vesicles, in which microscopically visible, liquid-ordered (L_o_) and liquid-disordered (L_d_) lipid domains form upon a shift to low temperatures. The extent such phase behavior exists in the membrane of the endoplasmic reticulum (ER) of cells remains unclear. To explore the phase behavior of the ER membrane in cells, we used hypotonic cell swelling to generate Large Intra-Cellular Vesicles (LICVs) from the ER in cells. ER LICVs retained their lumenal protein content, could be retubulated into an ER network, and maintained stable inter-organelle contacts, where protein tethers are concentrated at these contacts. Notably, upon temperature reduction, ER LICVs underwent reversible phase separation into microscopically-visible L_o_ and L_d_ lipid domains. The L_o_ lipid domains marked ER contact sites with other organelles. These findings demonstrate that LICVs provide an important model system for studying the biophysical properties of intracellular organelles in cells.

**Significance Statement:** Prior work has demonstrated that the plasma membrane can phase separate into microscopically visible L_o_ and L_d_ domains with distinct lipid and protein content. However, such behavior on the ER membrane has not been experimentally observed, even though the ER contacts every organelle of the cell, exchanging lipids and metabolites in a highly regulated manner at these contacts. We find here that hypotonic treatment generates Large Intra-Cellular Vesicles from the endoplasmic reticulum and other membrane-bound organelles in cells, enabling the study of phase behavior on the ER membrane. We show that ER membranes can be reversibly phase separated into microscopically-observable, L_o_ and L_d_ domains. ER LICVs also maintained stable inter-organelle contact sites in cells, with organelle tethers concentrated at these contacts.

## Introduction

It has long been hypothesized that small, diffraction-limited, liquid-ordered micro-domains can form on the plasma membrane of cells^1–3^. Lipids such as cholesterol and sphingolipids are concentrated in these domains, and membrane and membrane-associated proteins can preferentially segregate into these domains to allow sorting of proteins and localized signal transduction^2,4,5^. An important tool for studying these properties of the plasma membrane (PM) has been giant plasma membrane vesicles (GPMVs), which are large vesicles that bud off from the plasma membrane of adherent cells under certain conditions ^6–9^. GPMVs contain the complex mixture of lipids and proteins found in the intact PM and are not contaminated with membranes from other organelles^9^. GPMVs can phase separate into microscopically-visible L_o_ and L_d_ lipid domains that have distinct sets of proteins and lipids, making them an important model system for studying the domain preference of membrane proteins^8,10,11^. Similar phase-like, membrane heterogeneities are thought to occur in the intact PM, albeit on a smaller scale due to constant lipid turnover and membrane trafficking^12^.

Up until now, the use of GPMVs to study properties of cellular membranes and membrane proteins has been limited to membrane proteins and lipids that localize to the PM naturally or which can be forced to localize to the plasma membrane of cells^13^. However, many fundamental processes of protein sorting and signal transduction occur intracellularly, through the activity of membrane-bound organelles^14–16^. The various intracellular organelles (including ER, lysosomes, endosomes, peroxisomes, mitochondria and lipid droplets (LDs)) have distinct membrane compositions and communicate with each other through membrane trafficking pathways or through membrane contact sites, which are domains where membranes of two organelles are kept in close proximity by protein-protein or protein-lipid tethering complexes^17^.

Contact sites represent only a small proportion of the overall surface area of an organelle, but they serve several fundamental functions, including being the sites of inter-organelle exchange of lipids, ions, ROS and other small molecules ^17–21^. Contact sites also play an important role in organelle inheritance and fission ^17,20–28^. Even though known tethering, effector, and regulatory proteins involved in contact site organization have been uncovered, how contact sites assemble and whether they exhibit self-organizational properties of lipids and proteins is unclear^17^. This ambiguity is largely due to the difficulty of studying the time-dependent behavior of lipids and proteins at nanoscopic contact sites^9,12^.

Here, we demonstrate that hypotonic swelling of adherent cells forms ER-derived, large intra-cellular vesicles (LICVs) that can be used as a tool for investigating the self-organizing and phase behavior of ER membranes. ER LICVs do not rupture in cells and they can be rapidly tubulated to form a tubular ER network in cells. When cells are chilled below room temperature, ER LICVs display reversible, microscopically observable phase separation into L_o_ and L_d_ domains. Hypotonic swelling also generates stable LICVs from other membrane-bound organelles within cells and stable inter-organelle contacts between LICVs are maintained in cells. Inter-organelle tethers are stably concentrated at these contacts between LICVs. In chilled cells, L_o_ domains in the ER LICV membrane formed at these contact sites with other organelles. LICVs generated with hypotonic swelling thus represent a model system which can be used to study the biochemical and biophysical properties of intra-cellular organelles in eukaryotic cells.

## Results

### Hypotonic swelling vesiculates the ER in adherent cells

The ER is the largest organelle within the cell and it is responsible for many cellular processes including the synthesis of proteins, lipids, and fatty acids^29^. COS7 cells were transiently transfected to co-express Sec61β-GFP to identify ER membranes and mCherry-KDEL to label the ER lumen^30,31^. Before swelling, the ER existed as a fine tubular network extending throughout the cytoplasm and surrounding the nuclear envelope (Figure 1A: control). In these cells, the ER lumen and membrane markers had overlapping distributions due the diffraction-limited size of ER tubules. After incubation at 37 C for 10 minutes in hypotonic LICV media to cause cell swelling, the ER’s fine tubular network was transformed into numerous Large Intra-Cellular Vesicles (LICVs) comprised of Sec61β-positive membranes surrounding a large lumen filled with mCherry-KDEL (Figure 1A: swollen). No dispersal of mCherry-KDEL into the cytoplasm was seen, indicating the ER LICVs did not rupture within the cell. Cell-to-cell heterogeneity in the degree and size of ER LICV generation was observed, yet many cells (∼30-40 %) exhibited complete and dramatic transformation of the tubular ER network into micron-scale LICVs (Figure 1A: swollen).

**Figure 1.**
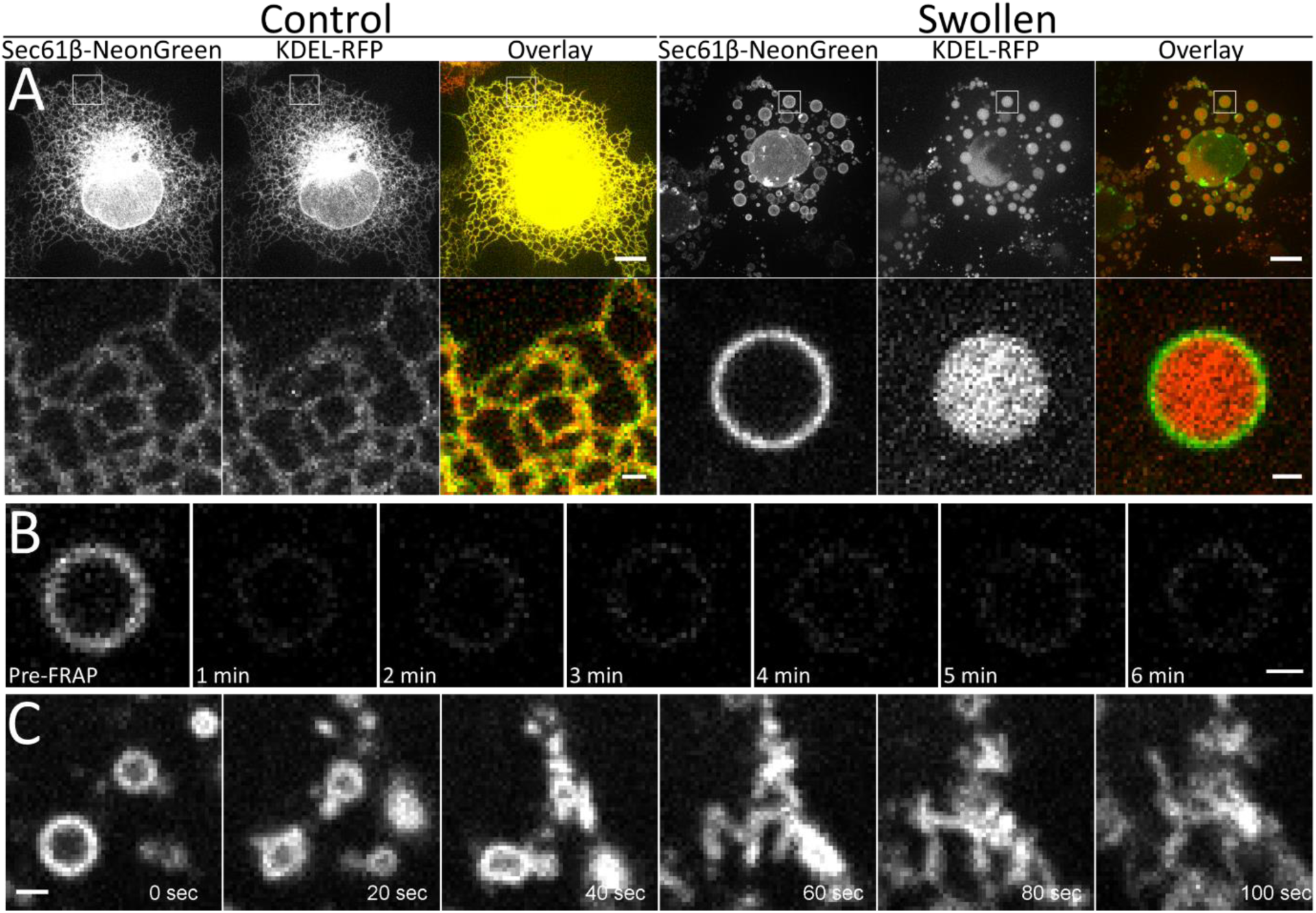
The effect of hypo-osmotic swelling on the ER of COS7 cells. **A.** Columns 1-3 show a representative cells in a proliferating that have been co-transfected to visualize the ER membrane and lumen with Sec61β-mNeonGreen and KDEL-RFP, respectively. Columns 4-6 show a representaive cell after hypotonic treatment. The second row is a scaled image of the region indicated by the square. Scale bars are 10 microns and 1 micron, respectively. **B.** ER LICV membrane fluorescence does not recover after photobleaching the entire vesicle. Sec61b-mNeonGreen fluorescence is shown before photobleaching and at 1 minute intervals afterward. Images are all shown with identical brightness and contrast settings and can be compared directly. Scale bar is one micron. **C.** ER LICVs retubulate into a reticular ER network within two minutes after cells are returned to full media. Scale bar is 1 micron. COS7 cells were imaged at 37C, 5% CO2.

Sec61β-mNeonGreen fluorescence associated with a single LICV did not recover after photobleaching (Figure 1B), indicating that there is no exchange of membrane proteins between separate ER LICVs. This suggests that these ER LICVs are independent entities within cells and are not connected via membrane linkages in cells. We often find that the lumenal space between inner and outer nuclear envelope became separated and filled with a large volume of ER lumen (Supplementary Figure 1A), suggesting displacement or disassembly of nuclear pores, which normally bridge inner and outer nuclear membranes. Consistent with this, imaging cells transfected with the nuclear pore component NUP50 linked to mEmerald revealed it was confined to areas of the nuclear envelope where the inner and outer membranes had not separated (Supplementary Figure 1B).

Given the drastic change in ER morphology from a tubular network to LICVs in swollen cells, we examined the ability to regenerate a tubular ER morphology in cells. LICV media was carefully aspirated and full media was rapidly applied to the cells while imaging the Sec61β-mNeonGreen ER membrane marker. ER LICVs rapidly tubulate and reconnect to form a reticular network of ER tubules within two minutes (Figure 1C and Supplementary Movie 1). These results suggest that the strategy of deswelling ER LICVS in cells can be used to study a variety of ER membrane phenomena in cells, including tubulation and fusion of ER membranes^[58]^.

### ER-LICVs exhibit reversible, temperature-dependent phase separation

The membrane of the ER is thinner than the PM due to its reduced cholesterol content and it has yet to be shown that this membrane has the physical-chemical capacity for temperature dependent phase-separation^9,16,32^. Thus, we investigated whether, like GPMVs, ER LICVs show temperature-dependent, phase separation into visible L_o_- and L_d_ lipid domains. We expressed two proteins with expected opposite L_o_- and L_d_-lipid phase domain preferences in the ER membrane. One protein was Sec61β-mCherry, an ER-resident membrane protein with a short 21 amino acid helical transmembrane domain that we expected to favor the L_d_ phase. The other protein was an ER-retained glycosylphosphatidylinositol (GPI) C-terminal anchored mNeonGreen (GPI-mNG), which is highly concentrated in cholesterol-enriched, L_o_ domains of phase-separated GPMVs^8,9,33^.

Examining ER LICVs at 37°C in COS7 cells revealed that both proteins were homogenously distributed across the ER membrane (Figure 2A). The radial intensity profile around the circumference of the vesicle is plotted for both constructs and shows homogenous fluorescence intensity distributions around the ER LICV. When ER LICVs are chilled to room temperature (∼23°C), diffraction-limited domains of GPI-mNG form on ER LICVs which are depleted in Sec61β (Figure 2B). The radial intensity profile of this vesicle reveals the anti-correlation of the Sec61β signal with the GPI anchor signal. An even more dramatic de-mixing of ER LICV membranes occurred when cells were chilled to less than 10°C, as shown in Figure 2C. At these temperatures, microscopically visible domains appeared with concentrated GPI-mNG and highly reduced Sec61β-mCherry localization (Figure 2C, radial intensity profile). These results show that ER LICVs undergo temperature-dependent phase separation. In phase-separated ER LICVs, L_d_ lipid domains enriched in Sec61β occupy most of the membrane, with L_o_ domains enriched with GPI-mNeonGreen seen in discrete spots.

**Figure 2.**
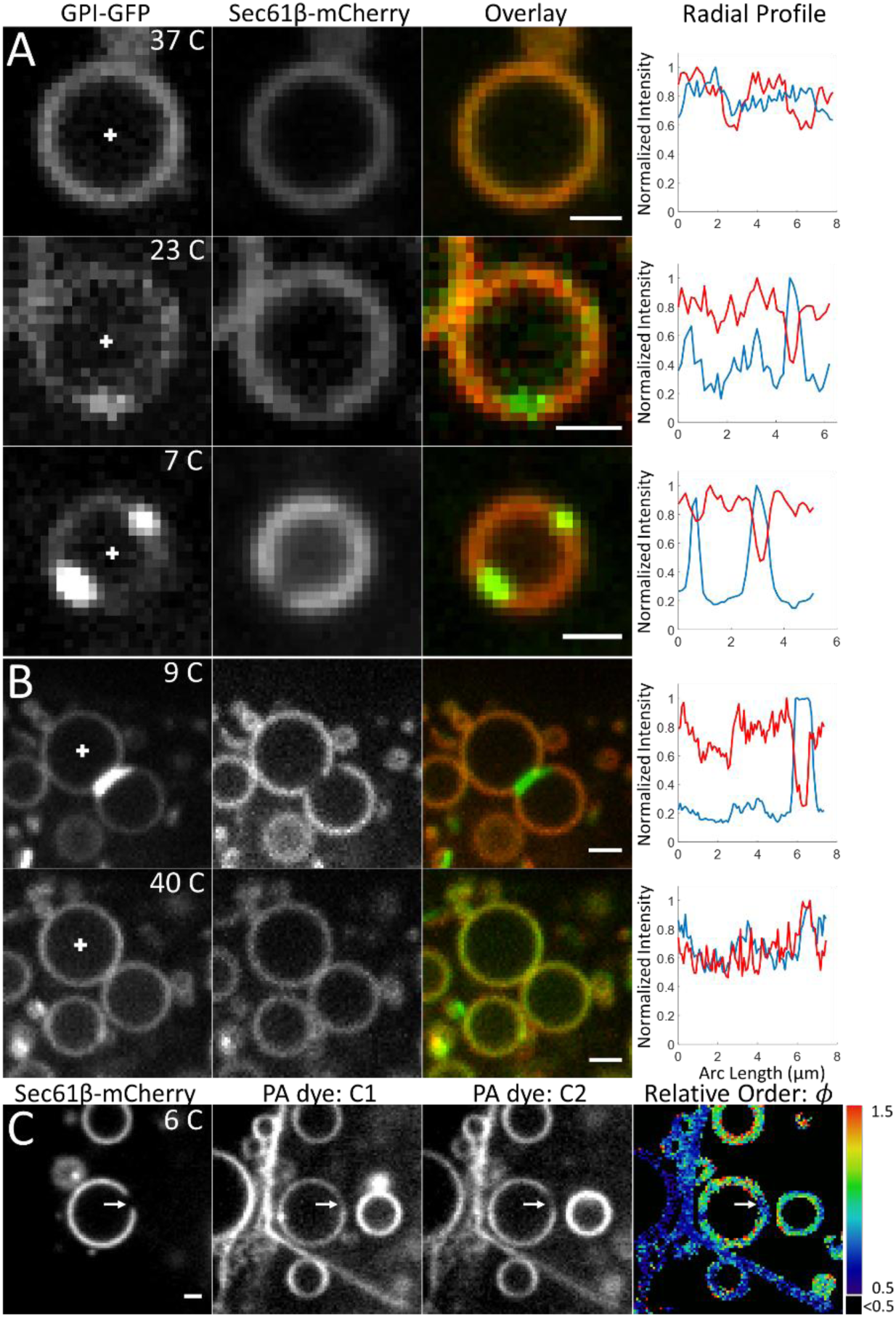
ER LICVs exhibit reversible, temperature-dependent phase separation. **A.** GPI-mNeonGreen was localized in the ER and colocalized with Sec61β-mCherry in ER LICVs at 37C. Microscopically observable domains begin to be observable at ∼23 C, with GPI-anchored mNeonGreen localizing to discrete puncta. Sec61β-mCherry fluorescence is suppressed at these locations. Chilling the cells to <10 C results in microscopically observable domains in ER-LICVs where the GPI anchor is localized to large puncta and Sec61b is excluded. **B.** Phase separated ER LICVs labeled with GPI-mNeonGreen and Sec61β-mCherry in a cell chilled to 9 ± 1 C. After quickly warming the cells to 40 C. domains that were once visible on the ER LICVs have disappeared. The temperature of the cells during imaging is indicated. Radial line profiles show the normalized pixel-level intensities for a circle drawn along the circumference of the vesicle, beginning at the x-axis and proceeding counter-clockwise. Sec61β signal is plotted in red and the GPI signal is plotted in blue. **C.** Ratio-metric imaging of the PA dye in phase separated ER vesicles shows a lower relative order parameter, *ϕ*, at regions excluding Sec61β. The colorbar indicates the fluorescence intensity ratio between the channels. Non-membrane-associated pixels have been segmented out from analysis and are displayed in black, along with all ratios < 0.5. Scale bars are 1 micron for **A-C**.

To test the reversibility of ER LICV phase separation, we induced phase separation in ER LICVs and then quickly warmed the warmed the cells. Figure 2B shows several phase-separated ER LICVs with large domains present on the membranes at 9 C. The radial intensity profile on the right panel of Figure 2B shows that the L_o_ domain on this vesicle is nearly a micron in length. Upon increasing the temperature to 40 C, the visible domains disappeared, and GPI-mNG and Sec61β-mCherry became uniformly distributed across the ER LICV membrane (Figure 2E and the line scan). Thus, temperature-dependent, phase separation of ER LIVCs into L_o_- and L_d_-lipid domains is reversible.

Independent confirmation of the temperature-dependent demixing of ER LICVs came with the use of a membrane order-sensitive, hydrophobic push-pull pyrene membrane dye, PA dye. PA dye labels all of the membranes of a cell and undergoes a spectral shift to shorter wavelengths in L_o_-domains, when compared to the spectrum emitted by the dye in L_d_ domains^34^. The dye fluorescence intensity is measured in GFP and RFP channels of the microscope and after correction for background, the relative order parameter, *ϕ* = *Int*_*RFP*_/*Int*_*GFP*_, is then lower in membranes with more order due to a spectral shift of the dye to shorter wavelengths. Figure 2C shows a phase-separated ER LICV that has been labeled with Sec61β-mCherry and the two channels of PA dye fluorescence. A large L_o_ domain excluding Sec61β is visible on the ER LICV (Figure 2C, left panel, arrow). The relative order parameter *ϕ* is also lower in this region of the ER LICV membrane indicating a spectral shift to the GFP channel (Figure 2C, right panel). This result confirms that the ER LICV membrane domains enriched in the GPI anchor and depleted in Sec61β are more ordered than the regions enriched in Sec61β on phase-separated ER LICVs.

### Hypotonic swelling generates Large Intra-Cellular Vesicles from other organelles

We next examined the effects of hypotonic swelling on the shapes of endosomes, lysosomes, mitochondria, peroxisomes and LDs (Figure 3). Endosomal membranes were labeled with GFP attached to two tandem FYVE (2FYVE-GFP) domains of the early endosome autoantigen 1 (EEA1) protein, which binds strongly to phosphatidylinositol 3-phosphate (PI3P), a phospholipid enriched in early endosomes ^35^. To label endosomal lumenal contents, cells were pulsed for 1-hour with AlexaFluor-594 labeled Dextran (10 kDa molecular weight)^36^. Prior to hypotonic swelling, endosomes were comprised mainly of small vesicles containing endocytosed dextran (Figure 3, Endosomes, Control). After swelling, these structures became enlarged, but did not release their luminal contents (Figure 3. Endosomes, Swollen). Small spherical structures excluding the dextran label were sometimes visible inside the swollen endosomes (see arrow). These are likely lumenal vesicles within the endosomal LICV.

**Figure 3.**
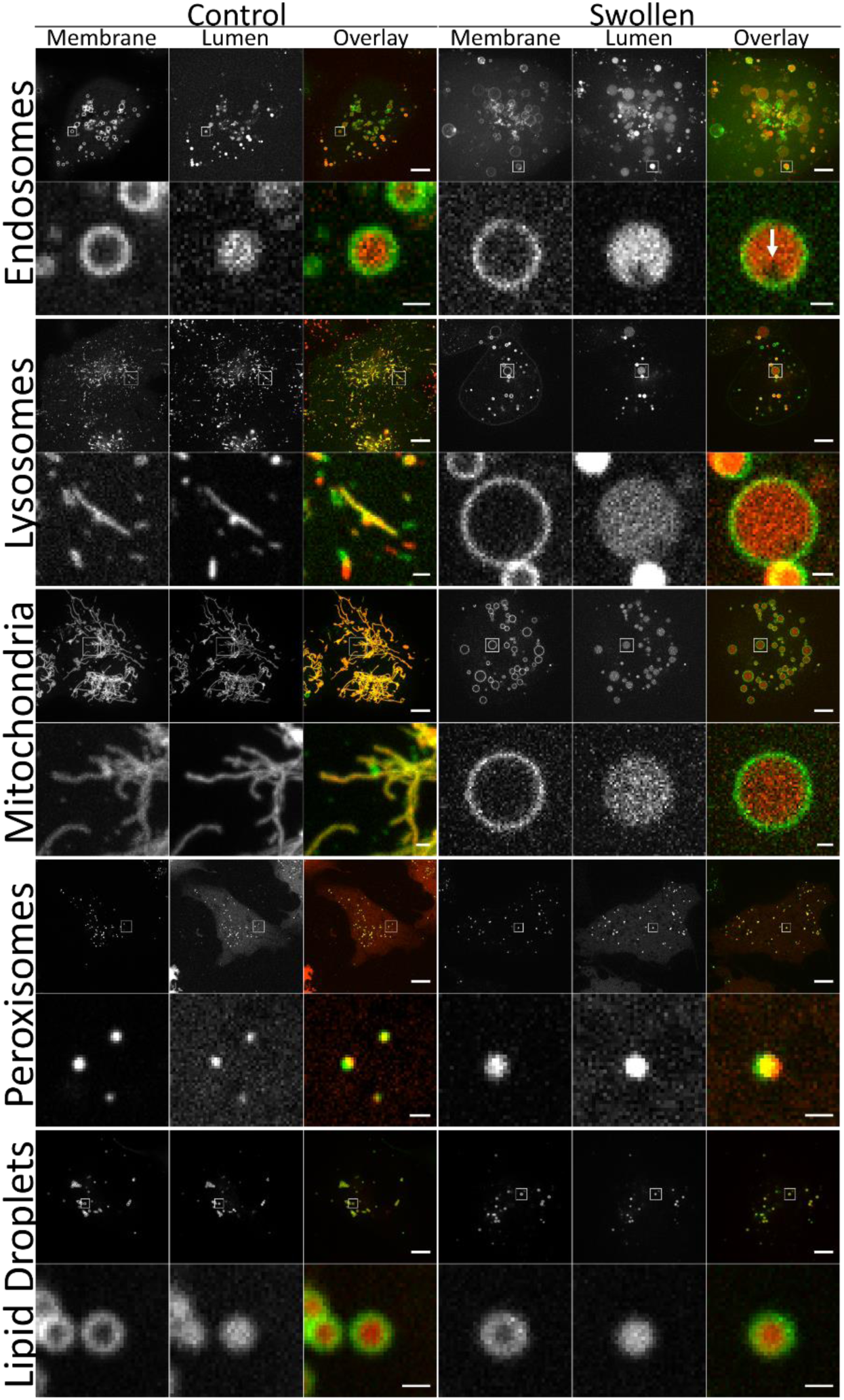
The effect of hypotonic treatment on the membrane bound internal organelles of COS7 cells. Columns 1-3 show a representative hypotonically swollen cell that has been co-transfected to visualize the the organelle membrane along with the lumenal contents. Columns 4-6 show a scaled region from the image in columns 1-3 indicated with the square. Endosomal membrane and lumen with 2FYVE-GFP, 10 kDa Dextran-AlexaFluor 594. Lysosomal membrane and lumen with Lamp1-YFP, 10 kDa Dextran-AlexaFluor 594. Mitochondrial outer membrane and matrix with TOM20-YFP, mTagRFP-mito. Peroxisomal membrane and lumen with PXMP2-mEmerald, SKL-mCherry. Lipid droplet phospholipid monolayer and neutral lipid core with ADRP-GFP, Bodipy-FarRed. Cells were imaged at 37C, 5% CO2. Scale bars are 10 micron and 1 micron, respectively.

To visualize the effect of hypotonic swelling on lysosomes, cells were transiently transfected with the lysosome-associated membrane glycoprotein (LAMP1)-YFP and treated with a 3-hour pulse-chase of fluorescent Dextran^36^. Before swelling, lysosomes were small vesicles or exhibited a tubular network morphology in cells (Figure 3, Lysosomes and Control). After swelling, lysosomes swelled into lysosomal LICVs with LAMP1-positive membranes surrounding a lumen filled with Dextran (Figure 3.Lysosomes, swollen). Some of the lysosomal LICVs had LAMP-positive vesicles inside of them, indicating they were possibly multivesicular bodies (Supplementary Figure 2).

Mitochondria were examined in cells co-labeled with the mitochondrial outer membrane protein TOM20 linked to YFP (TOM-YFP) and a mitochondrial matrix targeting sequence linked to mTagRFP (mTagRFP-mito)^37^. Hypotonic swelling drastically reshaped the tubular network of mitochondria (Figure 3, mitochondria, control) into numerous large, mitochondrial LICVs (Figure 3, Mitochondria, Swollen). In these vesicles, the matrix was swelled sufficiently to fill the space enclosed by the outer mitochondria membrane. The inner mitochondrial membrane is highly folded with cristae and can have a higher surface area than the outer membrane^29^. Consistent with this, herniated mitochondrial LICVs are also found in cells where the inner membrane and mitochondrial matrix extend into the cytosol from a large opening in the outer membrane (see Supplementary Figure 4). Thus, mitochondrial LICV generation exposes the difference in surface areas between the inner and outer mitochondrial membranes in cells.

**Figure 4.**
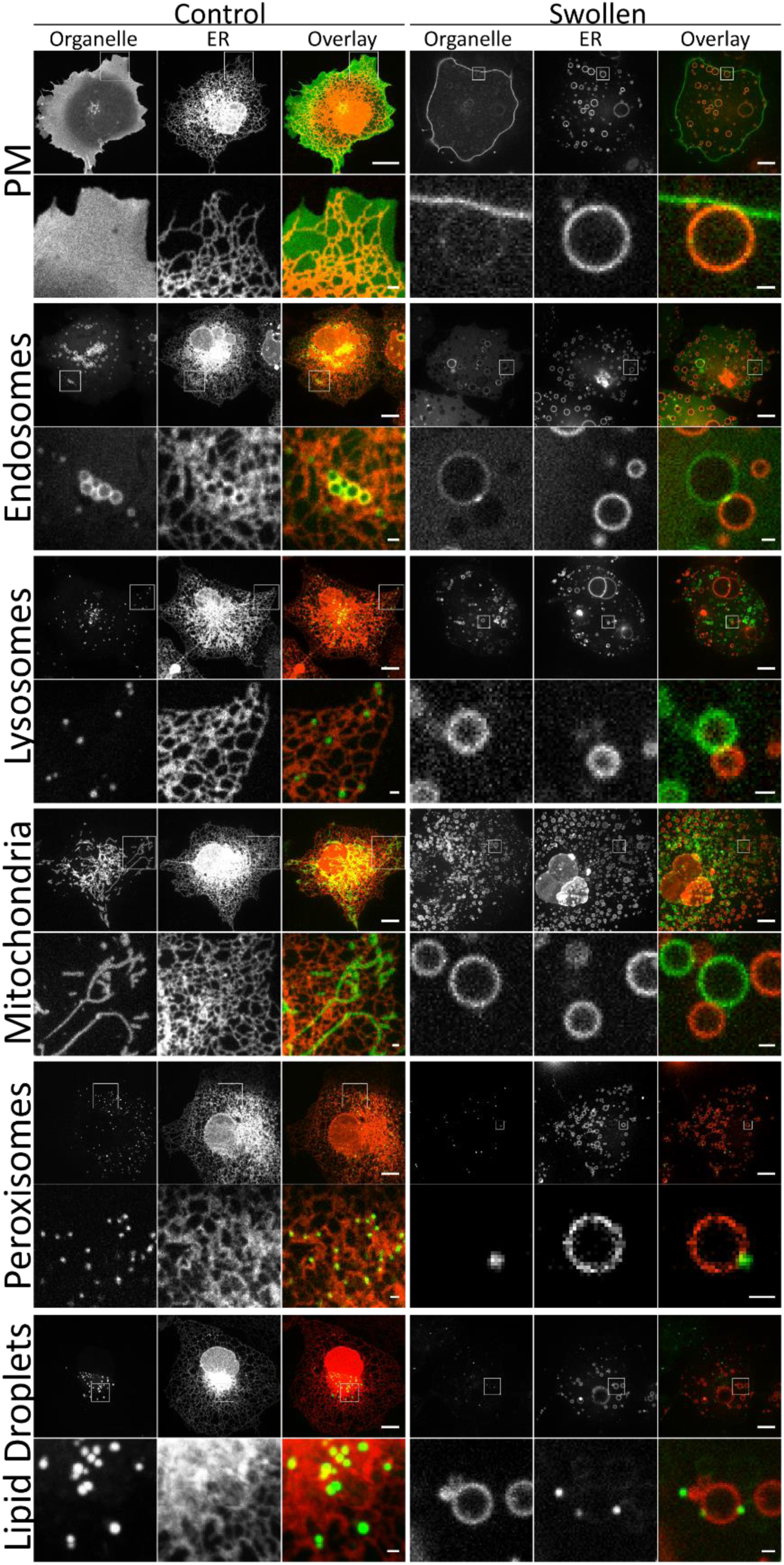
Contact sites with the ER are ambiguous in resting cells while contacts with ER LICVs are clear. Columns 1-3 show a representative, proliferating cell that has been co-transfected to visualize the ER along with the other organelles within the cell. Columns 4-6 hypotonically swollen cell. The region incdicated with a square is shown in a scaled version in the row below. The PM is labeled with GPI-GFP. Endosomes are labeled with 2FYVE-GFP. Lysosomes are labeled with LAMP1-YFP. Mitochondria are labeled with TOM20-YFP. Peroxisomes are labeled with PXMP2-mEmerald. Lipid droplets are labeled with Bodipy. The ER is shown in the second and fifth columns, labeled with Sec61β-mCherry, and shown in red in the overlays for consistency. The construct of columns 1 and 4 is shown in green in the overlays. Scale bars are 10 microns and 1 micron. COS7 cells were imaged at 37C, 5% CO2.

Peroxisomal outer membranes were labeled with the peroxisome-associated membrane protein 2 (PXMP2) fused to mEmerald^[31]^ and peroxisomal lumen was labeled with mCherry^38^. After treatment with LICV media, peroxisomes remained as diffraction-limited entities within cells and did not form LICVs (Figure 3. Peroxisomes, Control and Swollen).

The LD outer membrane was labeled through transient expression of adipose differentiation-related protein (ADRP) fused to GFP^39,40^. BODIPY 665/676 was then used to label the LD’s neutral lipid core. After hypotonic treatment, we found that the LDs retained their overall size and appearance in cells (Figure 3. LDs, Control and Swollen). Hypotonic swelling probably does not appreciably affect LD structure because the hydrophobic lipid core of the LD creates a phase-separated volume that already excludes water.

The overall state of cells after hypotonic swelling was examined with label-free, spatial light interference microscopy (SLIM, Phi Optics©). Swollen cells contained many LICVs, though the individual identity of these vesicles was not clear without a fluorescent tag (Supplementary Figure 4). Many LICVs were immobile within cells, while the other LICVs appeared to undergo random, thermal motion within the cell (see Supplementary Movie 2). Together, these results show that LICV formation occurs independently of the introduction of organelle-specific fluorescent tags and that hypotonic swelling generates LICVs from the ER, endosomes, lysosomes, and the mitochondria in cells.

### ER LICVs maintain stable inter-organelle contacts

Recent studies indicate that the ER establishes numerous contacts with the other organelles of the cell^17,26,29^. However, these contact sites are exceptionally difficult to study in live cells due to the complex, diffraction-limited, and time varying nature of inter-organelle contact sites. Thus, we investigated the possibility that inter-organelle contacts between LICVs were present in swollen cells. We first examined whether ER contacts with the PM, which serve as important sites for Ca^2+^ and lipid exchange, persist after hypotonic swelling^25^. Cells were co-labeled with the ER marker Sec61β-mCherry and the PM marker GPI-mNeonGreen^41^. After swelling, many ER-LICVs were found stably associated with the PM (Figure 4, PM). In this cell 39% out of 270 ER LICVs were associated with the PM.

Contact sites between the ER and endosomes were next examined. These contacts are known to coordinate endosomal fission and serve as important hubs for the transfer of cholesterol to and from late endosomes^24,26,42^. Cells were co-transfected with 2FYVE-GFP to label endosomes and Sec61β-mCherry to label the ER. After LICV media treatment, many endosomal LICVs were found stably tethered to ER LICVs, with a typical ER-endosome LICV pair (Figure 4, Endosomes). 90% of the 68 endosomes within this cell were found to be associated with ER LICVs. A clear concentration of the 2FYVE construct at the endosome-ER LICV contact site was also observed (see arrow pointing to punctum of concentrated 2FYVE-GFP), implying that PI3P is enriched at the endosomal-ER contact sites. Consistent with this, prior work has shown that protrudin, an ER-localized protein with a PI3P sensing FYVE domain is concentrated at ER-late endosome contact sites^43^. Conversely, we find that Rab5b-GFP, another endosomal marker, is not concentrated at ER-endosome LICV contact sites (Supplementary Figure 5)^44^.

**Figure 5.**
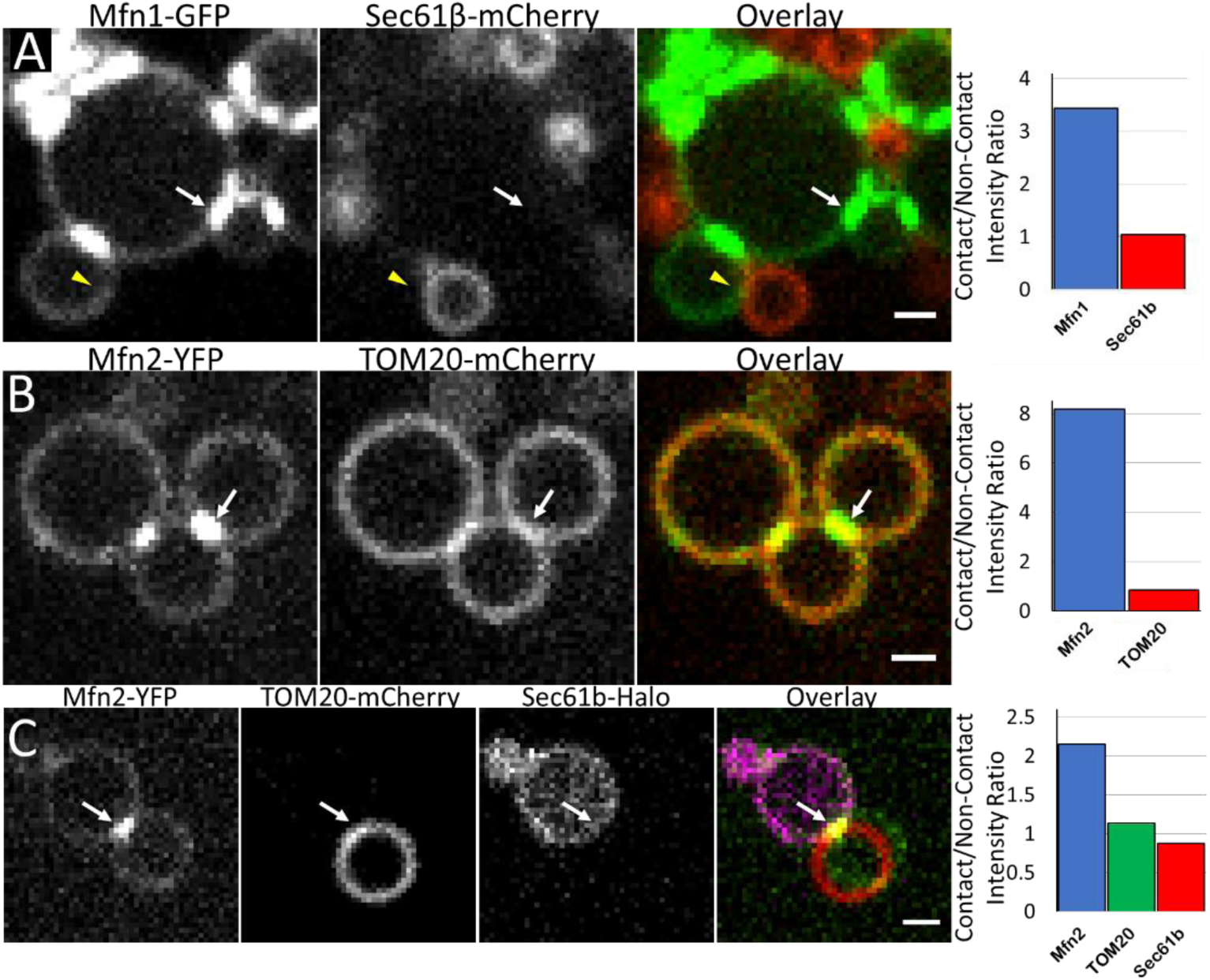
Tethers are concentrated at inter-LICV contact sites. **A.** Mfn1 localizes to mitochondrial LICVs and puncta of Mfn1 form at inter-mitochondrial contact sites. From left to right: Mfn1-GFP, Sec61β-mCherry, and overlay. The bar plot shows Mfn1(blue) enhancement at the mito-mito contact site indicated by the white arrow, but Sec61β is not enhanced at the ER-mito contact site (yellow carrot). **B.** Puncta of Mfn2 form at inter-mitochondrial LICV contact sites. From left to right: Mfn2-YFP, TOM20-mCherry, Overlay. The bar plot shows Mfn2 (blue) enhancement at the inter-LICV mitochondrial contact site, but TOM20 (red) is not concentrated at the contact indicated by the white arrow. **C.** Puncta of Mfn2 form at ER-mitochondrial LICV contact sites. From left to right: Mfn2-YFP, TOM20-mCherry, Sec61β-Halo labeled with JF646. Overlay. The bar plot shows Mfn2 (blue) enhancement at the ER-LICV mitochondrial contact site, but TOM20 (green) and Sec61β are not concentrated at the contact indicated by the white arrow. Scale bar is one micon for **A-C**. Cells were imaged at 37 C, 5% CO2.

ER-lysosome contacts are important sites for regulated transfer of Ca^2+^ from the ER to lysosomes^45^. To examine whether these contact sites persist after LICV media treatment, cells were co-transfected with LAMP1-YFP to visualize lysosomes and Sec61β-mCherry to label ER. After cell swelling, lysosomal LICVs were also found associated with ER LICVs in cells (Figure 4, Lysosomes). In this cell, 22% of the 590 lysosomal vesicles were associated with ER LICVs.

ER-mitochondrion contacts mediate lipid and Ca^+2^ transfer between ER and mitochondria ^29,46^. To see if stable ER-mitochondrion LICV contacts persisted in swollen cells, cells were co-transfected with TOM20-YFP to label mitochondria and Sec61β-mCherry to label the ER. After LICV media treatment, many stable contacts between ER LICVs and mitochondrial LICVs were found (Figure 4, Mitochondria). Of the 626 mitochondrial vesicles in this field of view, 96% were associated with ER LICVs.

The ER also forms contact sites with peroxisomes and LDs ^26,47^. ER-peroxisome contact sites are thought to be involved in the metabolism of long-chain fatty acids and ER-LD contacts are sites of transfer of fatty acids and phospholipids^27,48^. Cells were co-transfected with Sec61β-mCherry to label the ER and PXMP2-mEmerald to label peroxisomal membranes. After LICV media treatment, ER LICVs were found in cells with peroxisomes on their surfaces (Figure 4, Peroxisomes). In this cell, 93% of 267 peroxisomes were associated with ER vesicles. BODIPY 493/503 was then used to label the hydrophobic core of lipid droplets in cells transfected with Sec61β-mCherry to label the ER. After LICV media treatment, numerous LDs were also found on the surface of ER LICVs (Figure 4, LDs). 56% of the 206 lipid droplets in this cell were associated with ER vesicles.

Together, these results show that ER LICVs maintain contacts with all 5 major organelles in swollen cells. As the sites of contact between organels were clearly observable because of their enlarged volumes and the static nature, we could begin asking questions related to the biochemical properties of inter-organelle contact sites.

### Tethers are concentrated at inter-organelle contacts

Tethers are proteins that serve to functionally bridge opposing membranes of organelles to distances typically less than 40 nm, enabling inter-organelle communication and molecular transfer^17^. The primary requirements for a protein to be considered a tether are that it is concentrated at organelle contact sites and that it has a structural capacity to cause the close apposition of the two membranes through a protein-protein or protein-lipid interaction^17^. Given that numerous inter-LICV contacts occur within hypotonically swollen cells, we investigated the possibility that LICV generation can aid in the study of inter-organelle tethers.

The most well-characterized set of tethers in mammalian cells are the mitofusins: mitofusin 1 and mitofusin 2 (Mfn1 and Mfn2). Mfn1 and Mfn2 are mitochondrial outer membrane tethering proteins that are essential for mitochondrial fusion. Since Mfn2 has been shown to also localize to the ER, it has been proposed that it serves an additional role in ER-mitochondria tethering^22,28^. The mitofusins are enriched in cells at inter-mitochondrial contact sites through homotypic (Mfn1-Mfn1) or heterotypic (Mfn1-Mfn2) interactions^22,49,50^. To explore the distributions of mitofusins on LICVs at inter-LICV contact sites due to homotypic Mfn interactions, cells were transfected with Mfn1-GFP or Mfn2-YFP along with mitochondrial and ER membrane markers^51^ (Figure 5).

Figure 5A shows Mfn1-labeled mitochondrial LICVs in a cell co-transfected with the ER marker, Sec61β-mCherry. Clear puncta of Mfn1 are visible at mitochondrion-mitochondrion LICV contacts in this cell (white arrow), while puncta of Mfn1 do not appear at ER-mitochondrial LICV contacts (yellow carrot). The bar plot of Figure 5A shows that Mfn1-GFP fluorescence is highly enriched at inter-mitochondrial LICV contacts, but not with ER-mitochondrial LICV contacts.

As Mfn2 can localize both to mitochondria and the ER within cells, we performed separate experiments to examine Mfn2 at inter-mitochondrial and ER-mitochondrion LICV contacts^22^. Figure 5B shows several mitochondrial LICVs labeled with Mfn2-YFP and TOM20-mCherry. As with Mfn1, the bar plot of Figure 5B shows that Mfn2 (blue bar) is concentrated at inter-mitochondrial LICV contacts, while TOM20 (red bar), which has no tethering function, is not concentrated at the contact site. A small fraction of transfected COS7 cells showed Mfn2 localization on both ER and mitochondrial membranes. Thus, we triply transfected cells with Mfn2-YFP and TOM20-mCherry to label the mitochondria, and Sec61β-Janelia Fluor^®^ 646 to label the ER. Figure 5C shows an ER LICV and a mitochondrial LICV with Mfn2 localized to the membrane both vesicles. While Mfn2 is concentrated at the ER-mitochondria LICV contact site, TOM20 and Sec61β are not concentrated at the LICV contact site (Figure 5C, bar plot). This result provides independent support for the ER-mitochondrial tethering function of Mfn2, first proposed by de Brito and Scorrano^22^. Collectively, these results provide a straightforward confirmation of the known tethering properties of the mitofusins in cells.

### Phase separation of ER LICVs occurs at inter-organelle contact sites

Finally, we investigated the relationship between the ER-organelle contact sites and the locations of phase-separated L_o_-like and L_d_ domains found on chilled ER LICVs. As Sec61β-mCherry was enriched in L_d_ domains and was depleted from L_o_-like domains of ER LICVs at low temperatures (see Figure 2), we used its distribution as a proxy for assessing L_o_-like and L_d_ domain formation on ER LICVs at the locations of ER contacts with other organelles (Figure 4).

We began by looking at ER contacts with PM after LICV treatment in chilled cells. We find that the region of ER-LICVs that contact the PM are depleted in Sec61β-mCherry (Figure 6A, PM). Thus, the ER-PM contact site is also the site of L_o_ domain formation on the phase-separated ER LICV membrane. We turned next to ER contacts with endosomes (Figure 6A, Endosomes). In phase separated ER LICVs, Sec61β-mCherry was depleted from the ER membrane at the ER-endosome contact site. Much like the PM, L_o_ domain formation on ER LICVs occured at ER-endosome contacts between LICVs in chilled cells. In contrast to the behavior at PM and endosome contacts, ER-lysosome contacts occurred at L_d_ lipid domains on ER LICVs, where the Sec61β signal intensity was high (Figure 6A, Lysosomes). ER-mitochondrial LICV contact sites were locations of L_o_ domains on phase-separated ER LICVs (Figure 6A, Mitochondria). Turning next to the non-LICV forming organelles, we found that peroxisomes are localized to the Sec61β-enriched, L_d_ lipid domains, with peroxisomes located at the edge of L_o_ domains in the ER membrane (Figure 6A, Peroxisomes). Looking finally at ER-LD contacts, we found that LDs are associated with the L_o_ domains on phase-separated ER LICV membranes (Figure 6A, LD).

**Figure 6.**
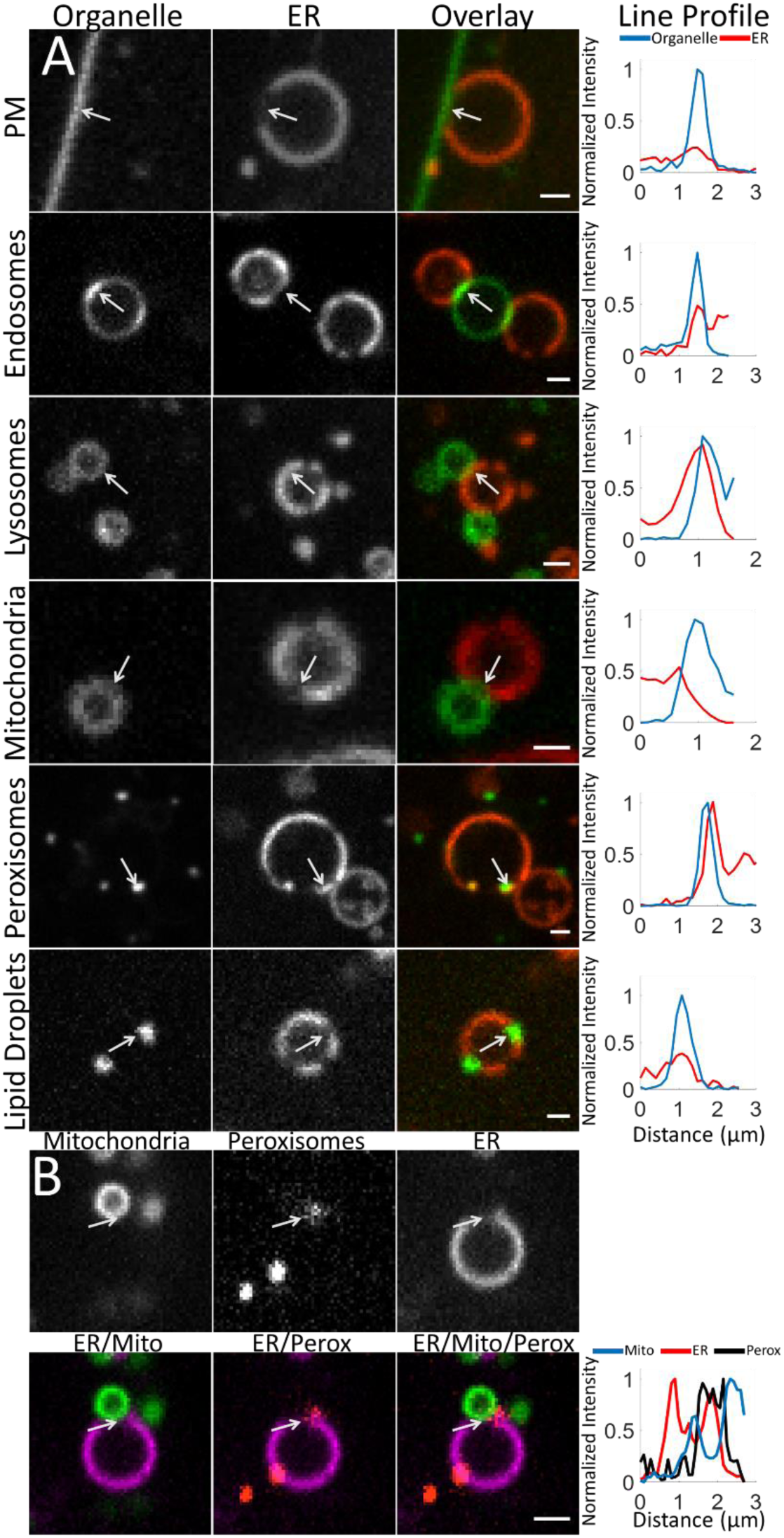
ER LICVs demonstrate phase behavior at inter-organelle contact sites. **A.** ER-organelle LICV pairs in cells that have been chilled to < 10 C. The plasma membrane is labeled with GPI-GFP. Endosomes are labeled with 2FYVE-GFP. Lysosomes are labeled with LAMP1-YFP. Mitochondria are labeled with TOM20-YFP. Peroxisomes are labeled with PXMP2-mEmerald. Lipid droplets labeled with Bodipy. The ER is shown in the second column in cells expressing Sec61β-mCherry, and is shown in red in the overlays. The construct of column 1 shown in green in the overlays. In the line profile, the organelle indicated by the row is plotted in blue and the ER is plotted in red. **B.** Three-color imaging single channel images and overlays of a phase separated ER-LICV labeled with Sec61β-Janelia Fluor^®^ 646, mitochondria labeled with TOM20-YFP, and peroxisomes with PXMP2-mCherry. Line profiles show normalized pixel-level intensities of a line drawn from the end of the arrow and across the contact site along the arrow. Normalization for the ER membranes is respect to max Sec61β fluorescence at a non-contact region, and for the organelle, normalization is with respect to the max at the contact site. Scale bars are 1 micron for **A** and **B**.

Based on the above results, there appeared to be a pattern in the preference for L_o_- or L_d_-domainlocalization within ER-organelle LICV contact sites. While lysosomes receive their membrane components from endosomes, the PM, endosomes, mitochondria, and lipid droplets are organelles with known lipid transfer functions with the ER. These organelles all contacted the L_o_ domains on phase-separated ER LICVS. Peroxisomes were associated with L_d_ domains at the edges of L_o_ on phase-separated ER LICVs.

Peroxisomes receive membrane components from both the ER and mitochondria, thus we investigated the possibility that peroxisomes at the edges of L_o_ domains were associated with mitochondria as well as the ER. We used three-color fluorescence microscopy to examine the localizations of ER, mitochondria, and peroxisomes in chilled, swollen cells. We find that peroxisomes can be localized to tri-junction organelle arrangements which are associated with the L_o_ domains on phase-separated ER LICVs (Figure 6B). The line scan across the contact site shows that a peroxisome (arrow) is associated with a L_d_ domain (coincident PXMP2 signal with the Sec61β signal peak) at the edge of a L_o_ domain (suppressed Sec61β signal) on the ER LICV. The mitochondrion is bound at the L_o_ domain of this ER LICV (TOM20 peak at position of Sec61β minimum). There is strong overlap of the mitochondrial signal with the peroxisomal signal at this location as well. Thus, peroxisomes can be associated with L_o_ domains on ER membranes through a tri-junction arrangement involving a mitochondrion. Collectively, these results suggest that organelles with lipid transport functions to and from the ER are associated with L_o_ domains on phase separated ER LICVs.

## Discussion

Little attention has been paid to the internal membranes of the cell under hypotonic conditions, despite its use to study cellular mechanisms of volume regulation and in creating spherically shaped blebs of plasma membrane to study protein-protein interactions ^52,53^. We have shown here that hypotonic treatment generates LICVs from most of the membrane bound organelles within the cell: the ER, endosomes, lysosomes, and mitochondria. LICVs are not generated from peroxisomes or lipid droplets.

The spherical shape of LICVs makes them amenable to a wide range of existing biophysical measurement techniques used to study the properties of lipids and proteins in the plasma membrane. For example, the LICV provides an ideal surface for membrane protein diffusion measurements, which are decoupled from the effects of the time-varying, complex, diffraction-limited topology of organelles in the proliferating cell. We also see that rescued cells will rapidly tubulate and fuse previously independent ER LICVs into a reticulated ER network (Figure 1C and Supplementary Movie 1). Thus, ER LICV generation could provide a means to study ER tubulation and fusion by enabling an initial state of the ER that is devoid of a diffraction-limited, complex topology. After hypotonic treatment, ER LICVs maintain stable inter-organelle contacts, enabling studies of biophysical activity at these contacts (Figure 4). We have shown here that LICVs enabled a clear observation that mitofusin 1 and 2 concentrated at inter-organelle tethering sites (Figure 5).

ER LICVs can be reversibly phase-separated into microscopically-observable L_o_ and L_d_ domains in a temperature-dependent fashion (Figure 2). Domain formation occurs at inter-organelle contacts with the ER, as shown in Figure 6. Therefore, LICV generation has shown that the ER membrane is capable of phase separating into microscopically visible L_o_ and L_d_ domains, like GPMVs made from the PM. In ER LICVs, ordered domain formation occurs at organelle contacts which are sites of lipid transport. Interestingly, lysosomes and peroxisomes are tethered to L_d_ domains on the phase-separated ER membrane, yet peroxisomes can be associated with L_o_ domains through an additional interaction with a mitochondrion. We see that not all inter-organelle contact are sites of L_o_ domain formation on ER LICVs, indicating that these different ER-organelle contact sites have different molecular properties.

GPMVs have been a crucial model system to understand physical properties and sorting mechanisms of plasma membrane components, together with other *in vitro* membrane model systems. Except for the recent discoveries of microscale phase-partitioned domains in yeast vacuole membranes, no tool to assess membrane properties and functions of intra-cellular organelles exists for eukaryotic cells^4^. Collectively, this study shows that the LICVs generated with hypotonic treatment provide a valuable new tool for the study of a wide variety of biophysical phenomena on the internal membranes of the eukaryotic cell.

## Methods

### Plasmid Constructs

In order of presentation within the text: mCherry-Sec61-C-18(Sec61β-mCherry), was a gift from Michael Davidson (Addgene plasmid #55129). Sec61β-mNeongreen was created by replacing the mCherry fluorescent protein with the mNeonGreen fluorescent protein with the NEB HiFi DNA Assembly Master Mix(E2621), according to the manufacturer’s instructions^54^. mCherry-ER-3 (KDEL-mCherry, vAddgene plasmid #55041), mEmerald-Nup50-N-10 (Nup50-mEmerald, Addgene plasmid #54210), mEmerald-PMP-C-10 (PMP-mEmerald, Addgene plasmid #54235), mCherry-Peroxisomes-2 (peroxisome lumen marker, Addgene plasmid #54520) were gifts from Michael Davidson. pEGFP-C1-ADRP (ADRP-GFP) was a gift from Elina Ikonen (Addgene plasmid #87161). TOMM20-YFP was created by replacing mCherry with the YFP fluorescent protein. mCherry-TOMM20-N-10 (TOMM20-mCherry, Addgene plasmid # 55146), mTagRFP-T-Mito-7 (mTagRFP-mito, Addgene plasmid # 58023) were gifts from Michael Davidson. pGFP-Cytochrome C was a gift from Douglas Green (Addgene plasmid # 41181). GPI-mNeonGreen was created by replacing GFP with mNeonGreen. pCAG:GPI-GFP (GPI-GFP) was a gift from Anna-Katerina Hadjantonakis (Addgene plasmid # 32601). Mfn2-YFP was a gift from Richard Youle (Addgene plasmid # 28010). Human mitofusin 1-GFP was purchased from OriGene (#RG207184).

### Cell Culture and Organelle Labeling

COS7 cells, cultured in DMEM containing 10% FBS and supplemented with L-glutamine, were seeded in Matrigel-coated 35mm glass-bottom petri dishes (Mattek). Twenty-four hours after seeding, at approximately 70% confluency level, cells were transiently transfected using Lipofectamine 3000 and 200-800ng of the plasmids indicated, according to the manufacturer’s instructions. Cells were imaged 12-24 hours after transfection.

For the loaded endosome experiments, cells were seeded and transfected as above with the 2FYVE-GFP construct. Just prior to imaging, a 1 hour pulse with 0.5 mM Dextran (10 kDa Dextran conjugated to AlexaFluor 594, ThermoFisher, #D22913), followed by extensive rinses with PBS buffer was performed. Cells were then swollen with hypotonic media and imaged. For the Dextran-loaded lysosome experiments, cells were pulsed for three hours with 0.5 mM Dextran, followed by extensive rinses with PBS buffer, and a chase of three hours in full media. Cells were then swelled and imaged.

### Cell Swelling

Cells were swelled with 1:20::DMEM:H2O diluted media, incubated for 10 minutes to allow for ER-LICV generation, and then imaged. We find that swollen COS7 cells returned to full tonicity with full media will undergo apoptosis and die, probably due to the presence of herniated mitochondria. Swollen cells are stable in the dish for over an hour. After some time, cells detach from the dish and have presumably died. Detached cells were not imaged and are not utilized in this study.

### Cell Microscopy

Live-cell confocal imaging was performed using a customized Nikon TiE inverted microscope with a Yokogawa spinning-disk scan head (#CSU-X1, Yokogawa) and an Andor iXon EMCCD camera. Fluorescence was collected through standard filters using a 100× Plan-Apochromat 1.40 NA oil objective (Nikon). Cells were imaged incubated with a Tokai Hit stage top incubator at 37°C and 5% CO2. Chilled cells were imaged under atmospheric CO2. Sample temperature was monitored and maintained with a QE-1HC Heated/Cooled Quick Exchange Platform controlled by a CL-100 Single Channel Bipolar Temperature Controller from Warner Instruments.

For the PA dye experiments, cells transiently transfected with Sec61β-mCherry were swollen with 1:20 DMEM containing 200 nM PA dye. After incubation at 37 C for 10 minutes, cells were chilled to induce visible phase separation in the mCherry signal, and PA dye signal was imaged with 405 nm laser diode excitation. Data was acquired sequentially in two scans with 400 ms exposure time for both on the GFP and RFP channels on the Nikon described above. During acquisition, the Andor iXon EMCCD camera gain was set to 1. A region containing no membrane was averaged to determine the background contribution in each channel’s data. The background was subtracted and the intensity ratio of the RFP to the GFP channel pixel values was calculated.

Spatial light interference microscopy was performed with a Phi Optics© SLIM unit attached to a Zeiss inverted microscope with a Plan-Neofluar” 63x/1.3 Imm Corr Ph3 M27 water immersion objective with a Hammamatsu C11440 digital camera. Cells were imaged at 37 C with 5% CO2 using a Tokai Hit stage top incubator.

## Data Analysis

Line scans and radial intensity profiles were created using the line-scan function in FIJI and LICVs were counted manually with the built-in Cell Counter plugin^55,56^. Data was plotted with MATLAB.

## Acknowledgements

The authors declare no competing conflicts of interests.

We are grateful to Dr. Ilya Leventhal and Dr. William Prinz for their thoughts and opinions. We are also thankful to Dr. Yacheng Liao, Dr. Heejun Choi, Dr. Chris Obara, and Dr. Chi-Lun Chang for their numerous minipreps, BODIPY and JF dyes, Sec61β-HaloTag construct, and experimental advice.

C.K. discovered LICVs while investigating ER shaping proteins. C.K., A.S., P.S., and J.L.S. designed the experiments, C.K. performed the experiments and data analysis. C.K. wrote the manuscript. C.K. created the figures. C.K., A.S., P.S., and J.L.S. edited the manuscript into final form.

---

**Fig. S1.**
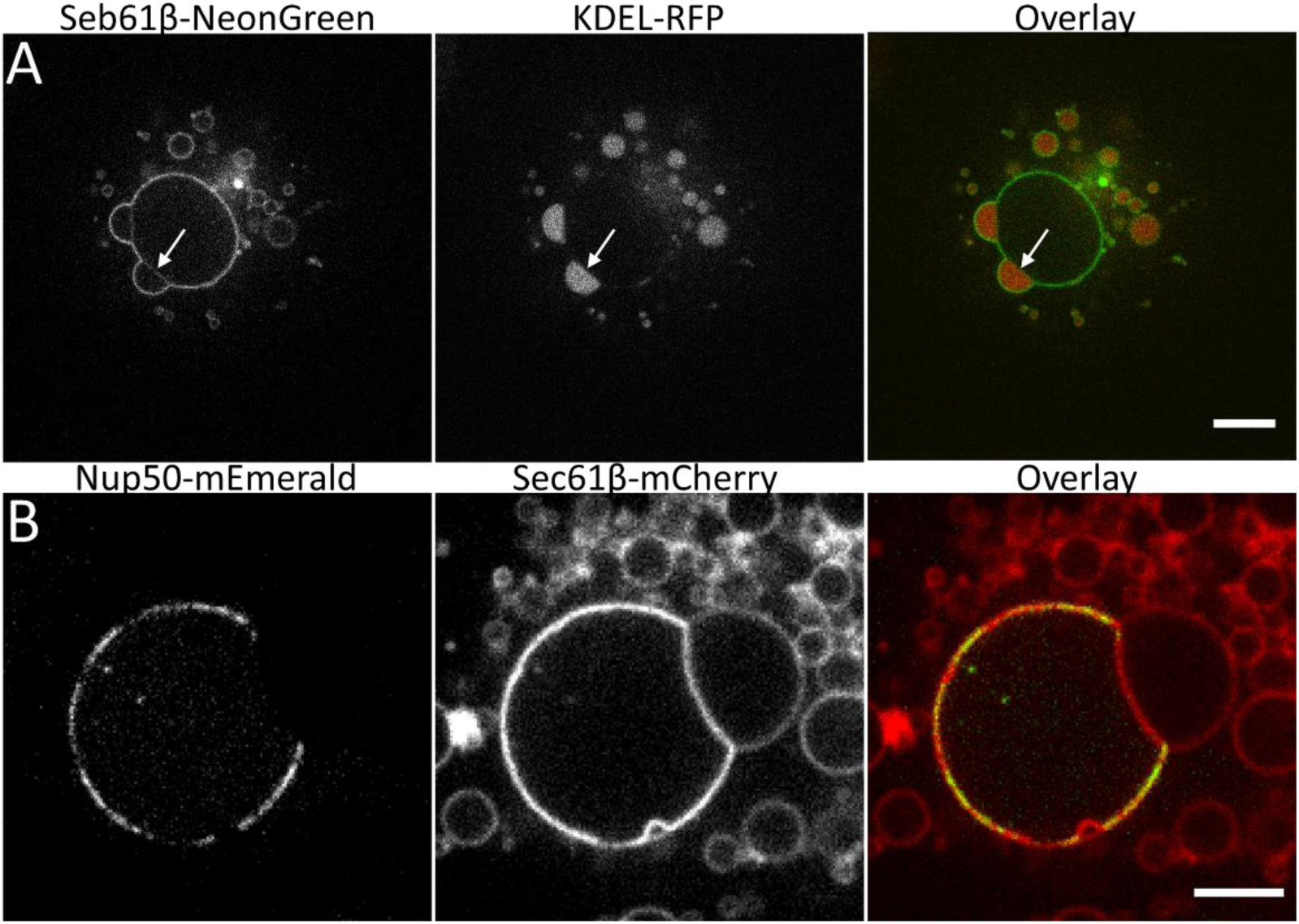
Nuclear membranes become separated after LICV media treatment. **A.** Large volumes of ER lumen are present on the nucleus in swollen cells where the inner and outer nuclear membranes are separately visible, as indicated by the arrow. Left: ER membrane labeled with Sec61β-mNeonGreen; center: ER lumen labeled with KDEL-mRFP; Right: overlay. **B.** Nuclear pores are excluded from the regions of separated inner and outer nuclear membranes. Left: Nup50-mEmerald; right: Sec61β-mCherry; right: overlay. Cells were imaged at 37 C, 5% CO2. Scale bars are 10 microns.

**Fig. S2.**
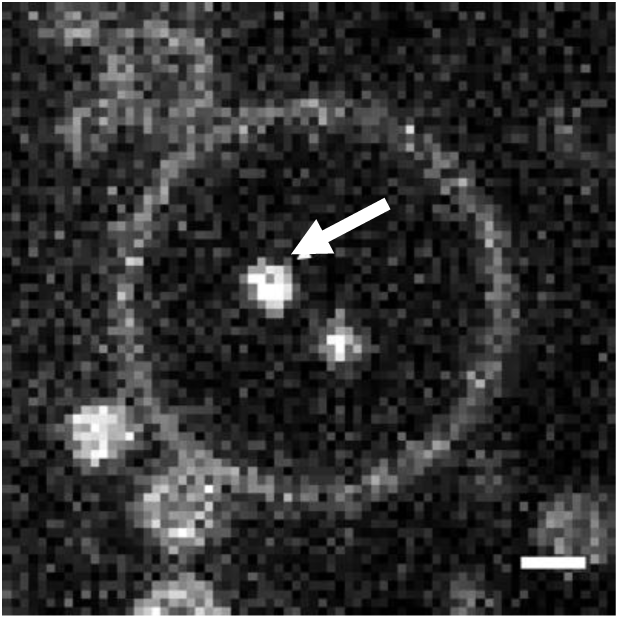
Small LAMP1-positive vesicles are present within lysosomal LICVs as indicated by the arrow. COS7 cell expressing LAMP1-YFP was imaged at 37 C, 5% CO2. Scale bar is 1 micron.

**Fig S3.**
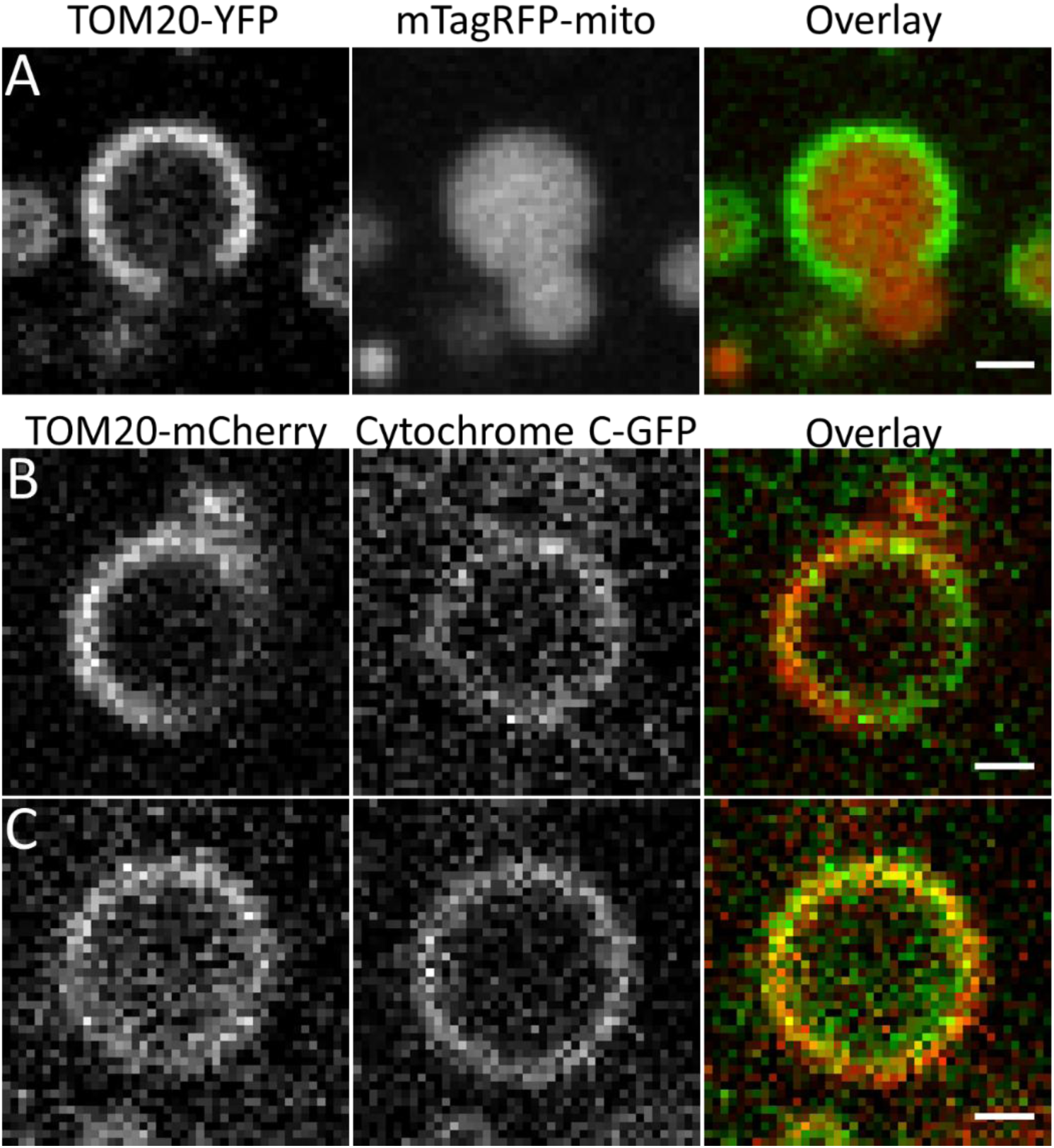
Hypotonic swelling herniates mitochondria. **B.** A mitochondrion with severe herniation. **C.** Herniated mitochondria display the inner mitochondrial membrane to the cytosol. **C.** A mitochondrial LICV with intact inner and outer membranes. Cells were imaged at 37 C, 5% CO2. Scale bars are 1 micron.

**Fig S4.**
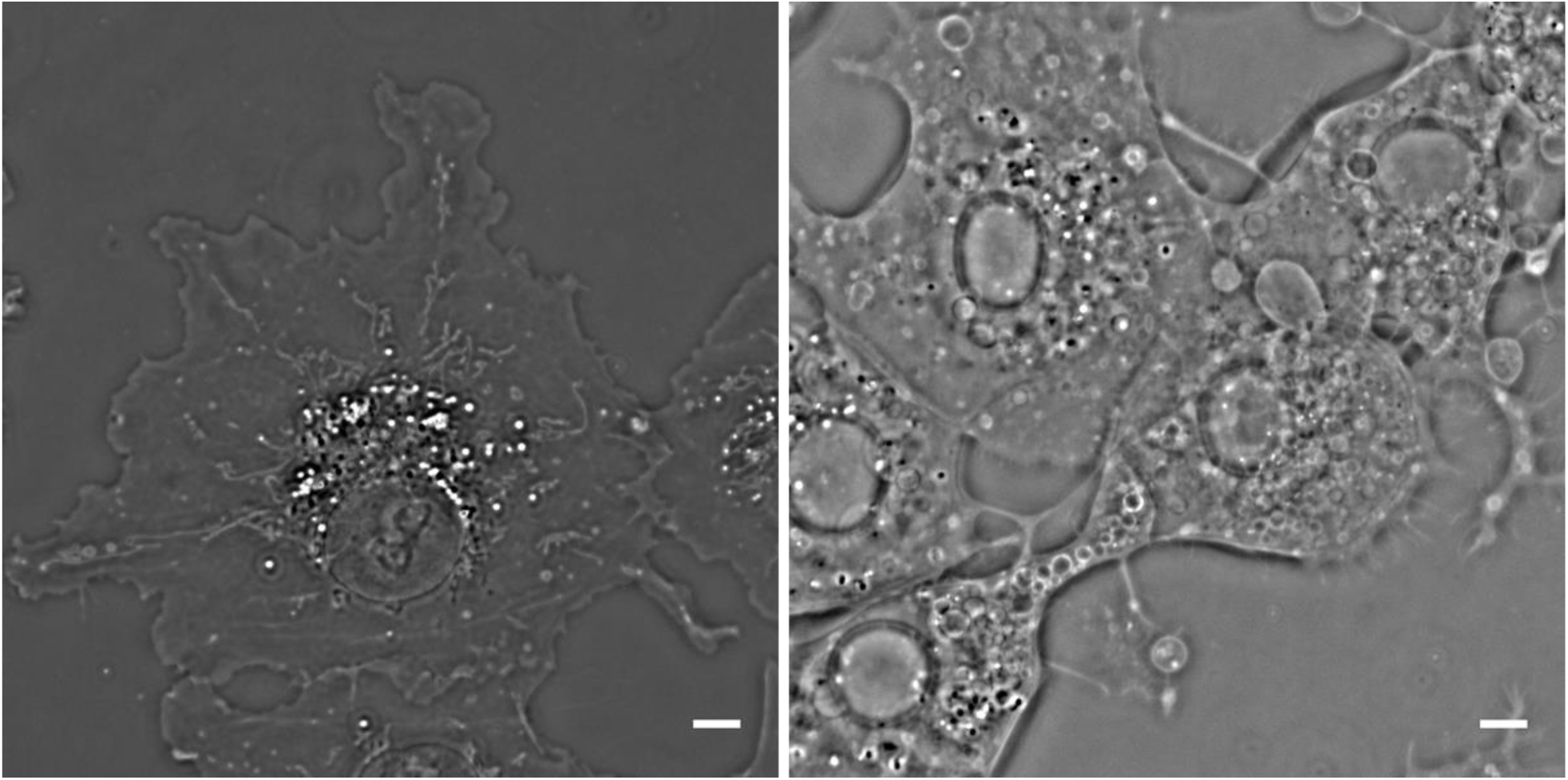
SLIM label-free imaging of the structure in resting and swollen cells. **A.** A resting COS7 cell. The nucleus, mitochondria, and other organelles are readily identifiable, and some regions of ER are also visible. **B.** Swollen COS7 cells contain with many LICVs from all the membrane bound organelles. COS7 cells were imaged at 37 C, 5% CO2. Scale bar is 10 microns.

**Fig S5.**
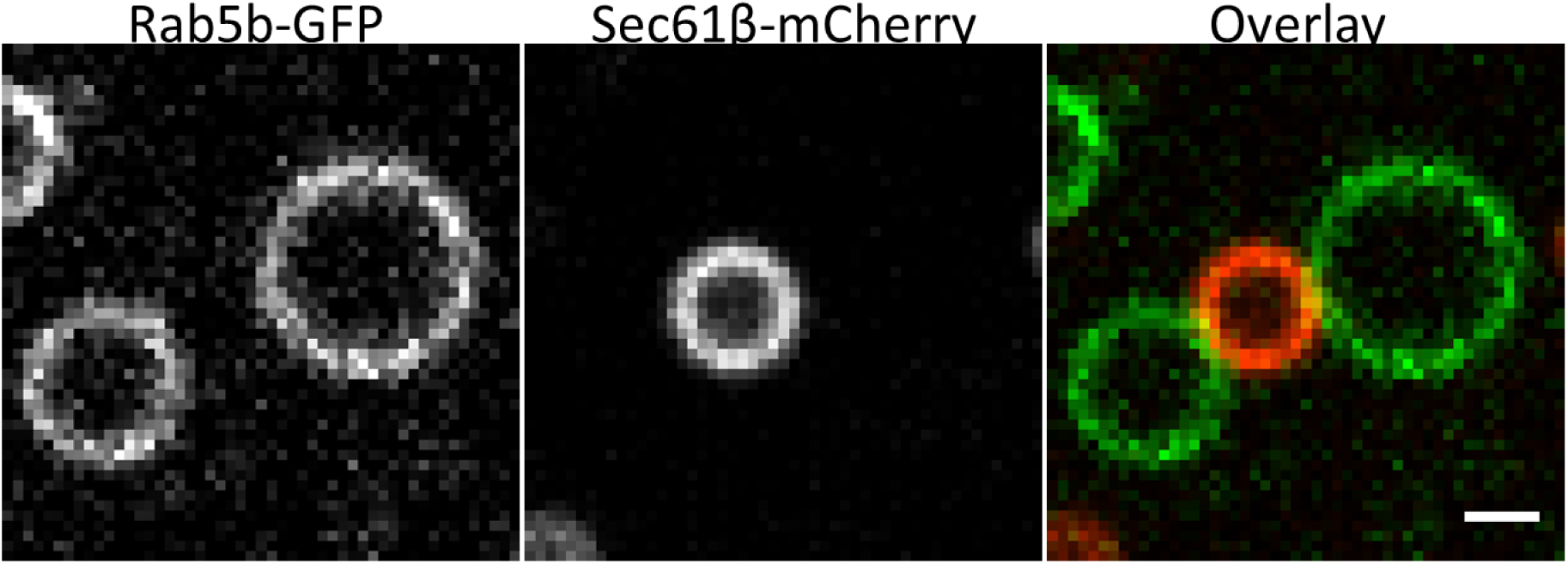
Rab5b is not enriched at ER-endosomal LICV contacts. Left: endosomes labeled with Rab5b-GFP Center: An ER LICV labeled with Sec61β-mCherry. Right: Overlay. COS7 cells were imaged at 37 C, 5% CO2. Scale bar is one micron.

**Movie S1.** Application of full media causes rapid retubulation of ER LICVs and the formation of an ER network in less than two minutes. Cells were imaged at 37 C, 5% CO2. Scale bar is one micron.

**Movie S2.** Many LICVs are immobile within cells, while the other LICVs undergo random motion within the cell. Movie shows a z-stack of COS7 cells after LICV media treatment, acquired with SLIM imaging at 1 micron steps. Cells were imaged at 37 C, 5% CO2. Scale bar is 50 microns.

